# Separations-encoded microparticles for single-cell western blotting

**DOI:** 10.1101/580233

**Authors:** Burcu Gumuscu, Amy Elizabeth Herr

**Author notes:** Footnotes relating to the title and/or authors should appear here. Electronic Supplementary Information (ESI) available: [details of any supplementary information available should be included here]. See DOI: 10.1039/x0xx00000x.

## Abstract

Direct measurement of proteins from single cells has been realized at the microscale using microfluidic channels, capillaries, and semi-enclosed microwell arrays. Although powerful, these formats are constrained, with the enclosed geometries proving cumbersome for multistage assays, including electrophoresis followed by immunoprobing. We introduce a hybrid microfluidic format that toggles between a planar microwell array and a suspension of microparticles. The planar array is stippled in a thin sheet of polyacrylamide gel, for efficient single-cell isolation and protein electrophoresis of hundreds-to-thousands of cells. Upon mechanical release, array elements become a suspension of separations-encoded microparticles for more efficient immunoprobing due to enhanced mass transfer. Dehydrating microparticles offer improved analytical sensitivity owing to in-gel concentration of fluorescence signal for high-throughput single-cell targeted proteomics.

## Introduction

The analysis of proteins in single cells plays a central role in identifying the link between cellular heterogeneity and disease states.[1,2] Like single-cell genomics and transcriptomics, single-cell protein analysis reports information that is concealed in bulk experiments.[3] For detection of protein targets known *a priori*, immunoassays are widely used in both analysis of large samples and to achieve single-cell resolution. Historically, heterogeneous immunoassays have been performed using a wide range of immobilizing substrates, including paper strips,[4] membranes,[5] and nozzle arrays.[6] Conventional immunoassays (e.g., ELISAs,[7] immunohistochemistry[8]) form the basis for the immobilizing substrates to study single-cell behavior, although specific protein types (such as protein isoforms that are crucial for cancer studies) cannot be detected in the absence of high specificity probes.[9] Other conventional immunoassays such as flow cytometry[10] and mass cytometry/CyTOF[11] can spatially resolve most protein isoforms, but intracellular protein targets are still difficult to measure in multiplexed runs and therefore analytical sensitivity remains insufficient for detection of key signaling proteins.[12] Macroscale immunoassays have been downscaled in order to improve sample detection and analysis capabilities by controlling mass and heat transport. At the microscale, heterogeneous immunoassays have used photopolymerized gel constructs,[13] microfluidic capillaries,[14] enclosed microfluidic channels,[15] microwell arrays,[16] and immuno-barcoding[17]. These approaches greatly simplified the target labeling process, facilitating a stepwise workflow of protein electrophoresis and antibody probing to discern off-target signals. However, a tradeoff still exists between maintaining satisfactory assay sensitivity and multiplexed analysis of > 1000 samples within the same batch that is required for measuring cell-to-cell variability in large populations.[18] Microparticles including barcoded hydrogel microparticles,[19] suspensions of particles,[20] and droplets [21] have also found utility in immunoassays; yet, the detection of target proteins directly from single cells could not be achieved in microparticle systems.

Miniaturization is well suited to electrophoretic separations owing to favorable scaling of physical phenomena including (1) efficient dissipation of Joule heating owing to high surface area to volume ratios found in microscale separation channels and (2) precision isolation and manipulation of individual cells (diameters ∼30 µm) – even among large populations of cells. For rapid electrophoretic analysis of single cell lysate, microchannel junctions and microwells prove useful for seamless handling of 1-5 pL of cell lysate. When an immunoassay is appended to a completed electrophoretic analysis, several additional advantages of miniaturization accrue. First, mass-based separation of proteins prior to an immunoassay separates any off-target, non-specific signal from that of the target [22] Second, controlled mass transport at the microscale shortens the probing and washout times of immunoassays. compared to surface-based immunoassay (e.g., microtiter plates). Suspended surfaces, such as microparticles, offer even more efficient mass transport owing to 3D access of reagents to the surfaces of the particle and, hence, reduced diffusion-length scales to the interior of the particle. Increasing the concentration of an immunoprobe can enhance immunoassay sensitivity via improved partitioning of immunoprobe into the particle.

Here, we introduce separations-encoded microparticles for single-cell immunoblotting, a hybrid approach that brings the selectivity of separations to the efficient compartmentalization of microparticles. The basis of the separation-encoded microparticles is a hydrogel molding and release technique, in which a planar array of microparticles is created with perforations delineating each microparticle perimeter. After use of the planar array for cell isolation and protein electrophoresis, the arrayed microparticles are mechanically released to create suspensions of microparticles, each encoded with a single-cell protein separation. We adopt the term “separations-encoded microparticles” to convey the concept that PAGE-separated proteins in each single-cell lysate are “coded” into the micron-size hydrogel, which is then released as a protein-patterned particle for subsequent immunoprobing and analysis. While the peak capacity of an electrophoresis separation scales inversely with separation lane length, the peak capacity of immunoblotting differs from that of pure separations, as the spectral channels afforded by the immunoprobing stage can allow targets that are not resolved by the electrophoresis be resolved by immunoprobing. By design, the hybrid device is designed for optimal performance at each assay stage. First, the planar hydrogel arrays are well-suited to sample preparation (i.e., isolation of a single cell in each microparticle using a microwell feature molded into each microparticle, in-microwell chemical cell lysis) and PAGE of single-cell lysate from the microwell into the abutting hydrogel and finally photoblotting-based protein target immobilization to hydrogel (Figure 1A, Figure 1B). Second, the microparticle form factor is well-suited to heterogeneous immunoassays (i.e., immunoprobing, washing) and further manipulation of hundreds-to-thousands of single-cell immunoblots. Dehydration of the microparticles shrinks the dimensions, yielding geometry-enhanced analytical sensitivity, with PAGE resolving power scale-invariant after blotting (immobilization). Lastly, we assess oncoprotein isoform expression across a range of cancer cell phenotypes. Separations-encoded microparticles bring performance benefits from both microarrays and microparticles to offer new avenues for high-throughput, high-selectivity protein cytometry.

**Figure 1.**
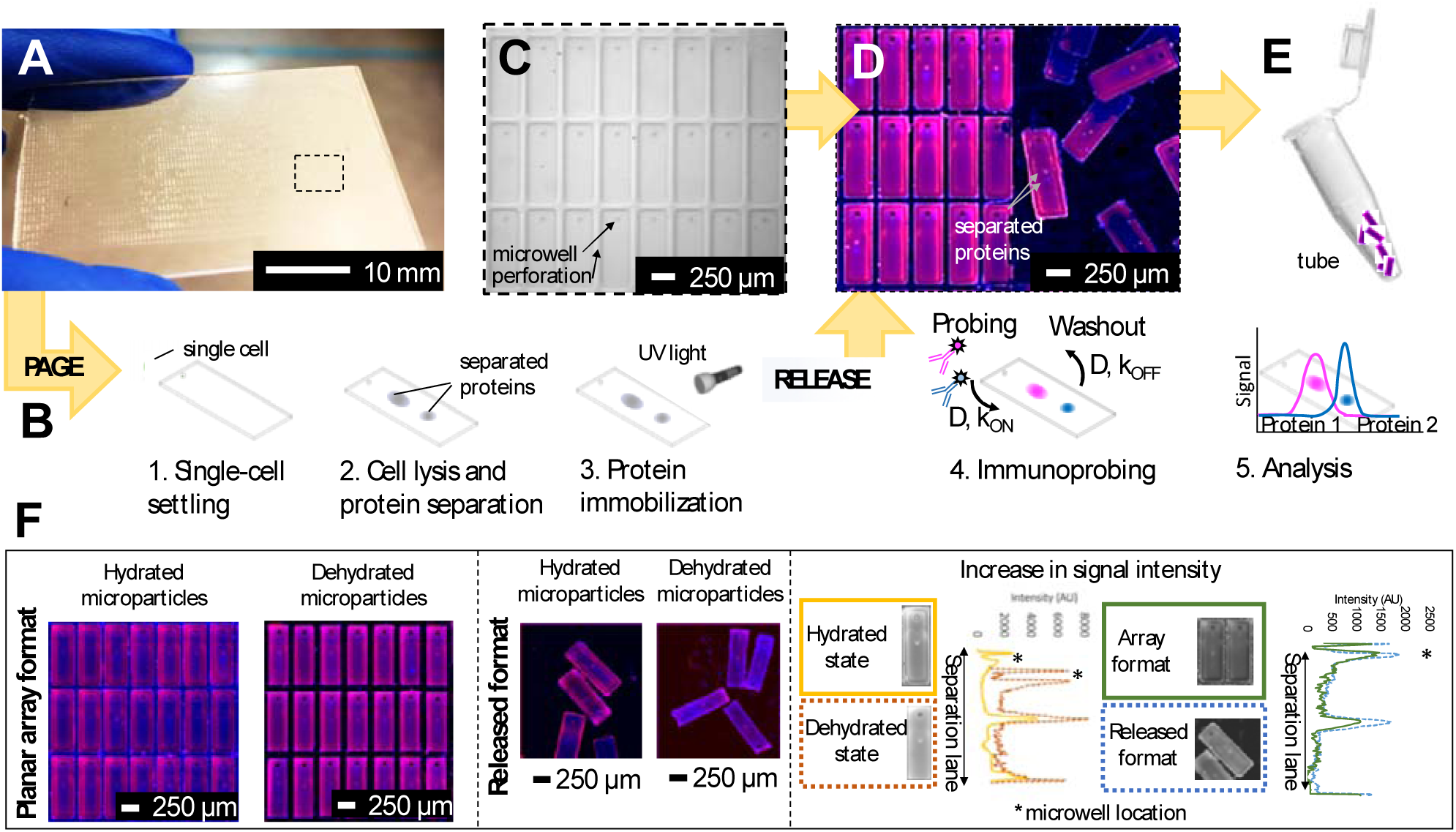
Design and operation of the separations-encoded microparticles for high specificity protein isoform analysis. **(A)** Separations-encoded microparticle array consisting of approximately 3500 releasable units fabricated on a half glass slide. **(B)** Schematic view of single-cell resolution western blotting workflow in microparticles. After performing the single-cell settling, electrophoresis, protein photocapture to the gel (immobilization), and immunoprobing, microparticles are released from the microscope glass slide by the help of a blade. Bands of separated proteins in each microparticle are visualized using a fluorescence scanner for quantification. **(C)** The array is comprised of a thin layer of polyacrylamide gel. Each microparticle contains a 30 µm diameter well where single cells are housed. **(D)** Single-cell resolution western blotting assay is ran on microparticles. Multiple protein markers can be probed in individual microparticles, the image shows false-colored micrographs of microparticles. **(E)** Microparticles can be released and collected for downstream analysis. **(F)** False-colored microscopy images of microparticle array attached on and released from a glass slide was probed and imaged in both hydrated and dehydrated states. Microparticles were probed for two housekeeping protein markers β-Tubulin (50 kDa, magenta) and GAPDH (35 kDa, blue) from single U251 cells. Dehydrating microparticles boosts analytical sensitivity thanks to the geometry-enhanced concentration of fluorescence signal, while the immunoprobe signal intensity in the released format is also increased attributable to enhanced surface area of each gel element.

## Results and discussion

### Lastly, Releasable separations-encoded microparticles: Single-cell immunoblots

Immunoblotting integrates PAGE of each single-cell lysate with a subsequent immunoassay to confer selectivity beyond PAGE alone.[18] In conventional protein immunoblotting, protein peaks are transferred from the PAGE hydrogel onto a hydrophobic membrane using electro-transfer or diffusion.[23] Hydrophobic interactions immobilize the protein separation on the blotting membrane, thus retaining separation information on a material with larger pore size than polyacrylamide molecular sieving gels. The larger pore size of blotting membranes (polyvinylidene difluoride (PVDF), nitrocellulose) reduces thermodynamic partitioning and enhances transport of immunoreagents to the immobilized protein material, thus underpinning efficient immunoprobing during the immunoassay.

We developed a method to create a planar array of releasable microparticles comprised of a dual-function hydrogel that, when comprising the planar array, acts as a molecular sieving matrix for electrophoresis and, when in a suspension of microparticles, acts as an immobilization scaffold for heterogeneous immunoassays performed on the separated protein targets. On silanized glass microscope slides (Figure S1A), we chemically polymerized polyacrylamide on an SU-8 mold to create planar microparticles, with perforations defining the perimeter of each microparticle. In selecting the microparticle shape, we sized each microparticle to house a microwell (15 µm radius, 40 µm depth) for mammalian cell isolation with an abutting region for protein PAGE. After single-cell PAGE analysis and photocapture of proteins to the hydrogel, the microparticles are released from the array by mechanical shearing, using a razor blade (Figure S1B). The limit of detection (LOD) of single-cell western blotting was previously characterized as ∼27,000 protein copies per cell, corresponding to the detection of the top 50% of most abundant proteins in the mammalian proteome [16]. This LOD value is determined by the signal acquisition technique (e.g., fluorescence microarray scanner), fluorophore-labelled antibodies, nonspecific background signal, and diffusive protein losses in assay steps.

To determine microparticle geometries, we sought to develop single-cell protein PAGE to analyze five ER-associated cancer signaling protein targets, spanning 35 kDa to 100 kDa (see Table 1) with a minimum mass difference between neighboring targets of 8% (i.e., β-tubulin and ERα46). The long axis of the rectangular microparticle (*L*_sep_) was determined using two separations-driven design criteria: (i) a target separation resolution (*SR*, defined as *SR* = *ΔL/4σ*, where *ΔL* is the separation length, *σ* is the average peak width) of 0.5 for the closest neighbors (*ΔL* = *ΔL*_min_) and (ii) the maximum electromigration distance (*L*_max_) for the protein target with the fastest electrophoretic mobility (*µ*, in-gel mobility, defined as *µ = µ*_0_ *10*^−*KT*^, where *K* is the retardation coefficient of an analyte and *T* is the total acrylamide concentration in the gel.[13,24] According to Ferguson analysis results, estimated electrophoretic mobility of proteins at 40 V cm^−1^ for 30 s in 8%T, 2.6%C gel is determined as shown in Table 1, (more details on electric field distribution in the array in SI)). Using these design rules, we fabricated planar arrays of rectangular microparticles ∼950-µm long, 250-µm wide, and 40-µm thick (Figure 1C). With the 950-µm long microparticles, we obtained a baseline resolution for intermediate molecular mass proteins with <30% mass difference, if immunoprobed together in the same probing cycle.[16] Our separations take the advantage that the separation lane length in immunoblotting scales directly with the peak capacity because immunoprobing step allows targets that are not resolved by the electrophoresis be resolved by immunoprobing. Upon mechanical release, we observed 94% yield of microparticles at hydrated state (n = 3 chips; 3500 microparticles per chip), with 91.4% of the successfully released microparticles exhibiting no discernable damage by visual inspection (Figure S2). Signal acquisition can be performed on the microparticle array or suspended microparticles (Figure 1D and Figure 1E), in either the hydrated or dehydrated state (Figure 1F).

**Table 1.** Protein targets in separations-encoded microparticles.

| Protein target | Molecular mass (kDa) | Mass difference between neighboring protein markers (%) | Electrophoretic mobility ($\text{m}^2 \text{V}^{-1} \text{s}^{-1}$) |
| --- | --- | --- | --- |
| GAPDH | 35 | 24 | $9.0 \times 10^{-9}$ |
| ERα46 | 46 | 8 | $6.8 \times 10^{-9}$ |
| β-tubulin | 50 | 25 | $6.0 \times 10^{-9}$ |
| ERα66 | 66 | 34 | $5.1 \times 10^{-9}$ |
| Actinin | 100 | - | $2.2 \times 10^{-9}$ |

We next validated the microparticle immunoblots through analysis of a range of breast cancer cell morphologies and types, using a panel of well-established breast cancer and kidney cell lines (breast adenocarcinoma, MCF 7; invasive breast adenocarcinoma, MDA MB 291; embryonic kidney cells, HEK 293). Average diameters of MCF 7, MDA MB 291, and HEK 293 cells were measured as 16 µm, 18 µm, and 16 µm, respectively; microwell diameters of 30 µm were used for all experiments. Based on the distribution of cell diameters in each cell population, cell settling resulted in zero, one, or multiple cells in microwells and perforations. In single-cell handling, we observed an average of 95% single-cell microwell occupancy for MCF 7, MDA MB 291 and HEK 293 cells, with 0.2% spurious isolation of cells in the perforations when a cell suspension of ∼ 10^6^ cells mL^−1^ in 1×PBS was introduced to the array (n = 4 devices, 3500 microwells per device), using a 10 min settling period (Figure S3A). A small number of microwells housed more than one cell (0.4%).

Two housekeeping protein targets, GAPDH and β-tubulin, were probed in the same microparticle assay using 10x diluted AlexaFluor 647 and AlexaFluor 555, respectively, and the resulting signal intensities were correlated with microwell occupancy (Figure S3B). Cells settled in the perforations did not have a detectable signal, which we attribute to rapid lysate dilution by convective flow (Figure S3B). After settling, in-microwell chemical lysis (30 s) and single-cell protein PAGE (20 s, *E* = 40 V/cm) were completed for GAPDH and β-tubulin across all cell types. Microparticle arrays were immunoprobed with a cocktail of GAPDH and β-tubulin (primary probing duration 3h, secondary probing duration 1h, washout periods 20 min; more details in Experimental Procedures). Observed electromigration behavior agreed with estimates (electromigration distances varying 0.9% and 1.1% for GAPDH and β-tubulin, respectively, 2 devices, n = 40 microparticles). With microparticles fabricated on silanized glass slides (Figure S4), electrophoretic mobilities of GAPDH and β-Tubulin were ∼9.0 × 10^−9^ m^2^ V^−1^ s^−1^ and ∼6.0 × 10^−9^ m^2^ V^−1^ s^−1^, respectively(Figure S3B), corroborated by previous observations [13] As expected, no protein signal was detected for empty microwells (n = 2 devices, 3500 microwells per device, Figure S3B).

In considering diffusive transport during immunoprobing, we hypothesized that immunoprobing of a suspension of microparticles would be more efficient than immunoprobing of the surface-attached planar microparticle array. The probing and washing steps dominate the planar assay duration. As context, conventional single-cell western blotting sees 75% assay duration devoted to probing and 25% to washout steps. Long durations are required owing to the limited mass-transport of antibody probes into the gel. In contrast, when in suspension the surface area of each microparticle is available for diffusion-based antibody probe introduction; whereas, transport into the hydrogel array is inhibited on the surface side that is attached to the glass slide. Then with 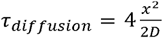 (where *τ*_*diffusion*_ is the transport time, *x* is the gel thickness, and *D* is the diffusion coefficient for antibody in an 8 %T gel [25]), *x* is the half-thickness of the microparticle thickness when the microparticle is in suspension and the full-thickness when the microparticle is anchored to a glass slide in the planar array. Figure 1F (right panel) compares immunoassay readout (fluorescence signal) from an immunoprobed protein separation performed in the array format and in the released format. We measure increased immunoassay signal upon in the released, particle format. We hypothesize that the increased immunoassay response derives from an increased surface area through which immunoprobe enters the bulk of the gel particle. The surface area is enhanced by releasing gel elements from the array format and suspending each gel particle in solution. The geometric argument applies to all stages of immunoprobing and washout, suggesting that the microparticle format could reduce the duration of the ∼4 hr immunoprobing-related steps by 25%. Regarding processing time estimates for one device (3500 particles): fluorescence imaging requires ∼ 30 min and signal analysis of micrographs (using MatLab) requires ∼15 min.

Background signal is also important to detection performance. Background is dictated by the efficacy of the washout process after probing. In comparing the planar array to the suspended microparticles, we observed background signal reduction of 1.3x in microparticles, and we indeed observed effective performance with reduced washout times (∼5 min vs. ∼20 min; n = 1 device, 3500 microparticles, CV = 0.2) in the suspension of microparticles versus the attached array (Figure S5A).

Next, in considering the immunoassay which is a reaction between a protein target and immunoprobe, as well as the transport during immunoprobing, we write *τ*_*reaction*_ = 1/(*k*_*on*_[*Ab*]_*gel*_ +k_*off*_) and [*Ab*]_*gel*_ = K[*Ab*]_0_, where *τ* is the reaction time, *k*_*on*_ is the reaction coefficient, *k*_*on*_ is forward reaction rate constant, *k*_*off*_ is backward reaction rate constant, *[Ab]*_*gel*_ is antibody concentration in gel, *[Ab]*_*0*_ is antibody concentration in solution—including partitioning coefficient for the hydrogel, *K* = 0.17 for 8%T PAG.(26) We use the Damköhler number (Da, with *Da* = *τ*_*ransport*_/*τ*_*reaction*_) [27] for low-affinity (K_D_ ∼10^−6^), medium-affinity (K_D_∼10^−9^), and high-affinity (K_D_ ∼10^−11^) immunoprobes.

Given this physical framework, we estimate that immunoprobing with low-affinity antibodies will be reaction-limited (Da∼0.7), while immunoprobing with medium-affinity (Da∼280) and high-affinity antibodies is mass-transport-limited (Da∼475).[26] From this analysis, we conclude that as long as medium- and high-affinity antibodies are used, the expedited diffusion into a 40-µm thick gel benefits the immunoprobing duration 4x faster in microparticle format when compared to the planar array format, according to *τ*_*diffusion*_ calculations.

We next sought to use the suspension of separations-encoded microparticles to overcome an important multiplexing limitation inherent to immunoblotting. In immunoprobing, an antibody pair is typically used to (i) detect the protein target (unlabeled primary antibody probe) and (ii) detect the unlabeled primary antibody (fluorescently labeled secondary antibody probe). The secondary antibody probe needs to be selective for the animal species in which the primary antibody probe was raised. Herein lies the detection challenge: primary antibodies are raised in just a handful of animal species. If multiple primary antibodies of the same species are used for target detection, the secondary antibody probes must be applied to the PAGE separation serially (not as one cocktail). The serial application demands multiple secondary antibody probing rounds and multiple gel stripping rounds, to ensure selective readout.[2] For example, two rounds of probing and stripping takes +50 hours for slab gel Westerns, and +9 hrs for conventional single-cell western blotting.[18]

To overcome this target multiplexing limitation, we fractionate the microparticle suspension into aliquots and apply distinct antibody probe solutions to each (i.e., Erα, Actinin). As a negative control, we performed two rounds of probing (for each probing round, 3 h primary and 1 h secondary antibody probing steps with 20 min washing time after the probing steps) and stripping (1 h) for ERα and Actinin antibodies separately (see Experimental Procedures). Figure 2A shows ERα expression level decreases in previously Actinin-probed microparticles, compared to microparticles probed for ERα alone. We calculated a 15.8% decrease in average expression quantified from negative control group (p>0.05, n = 40 microparticles). In multiplexed single-cell immunoassays, off-target probe binding is a substantial challenge (e.g., immunocytochemistry, flow cytometry).[7,8,10,11] Performing a separation, followed by immunoprobing helps to overcome this challenge by spatially separating the off-target signal. We investigated the off-target signals for both the ERα isoform and Actinin in separations-encoded microparticles and observed off-target signals for ERα (Figure 2B). Similar to slab-gel western blotting, signal is classified as off-target if detected outside the calculated target peak location (based on mobility).

**Figure 2.**
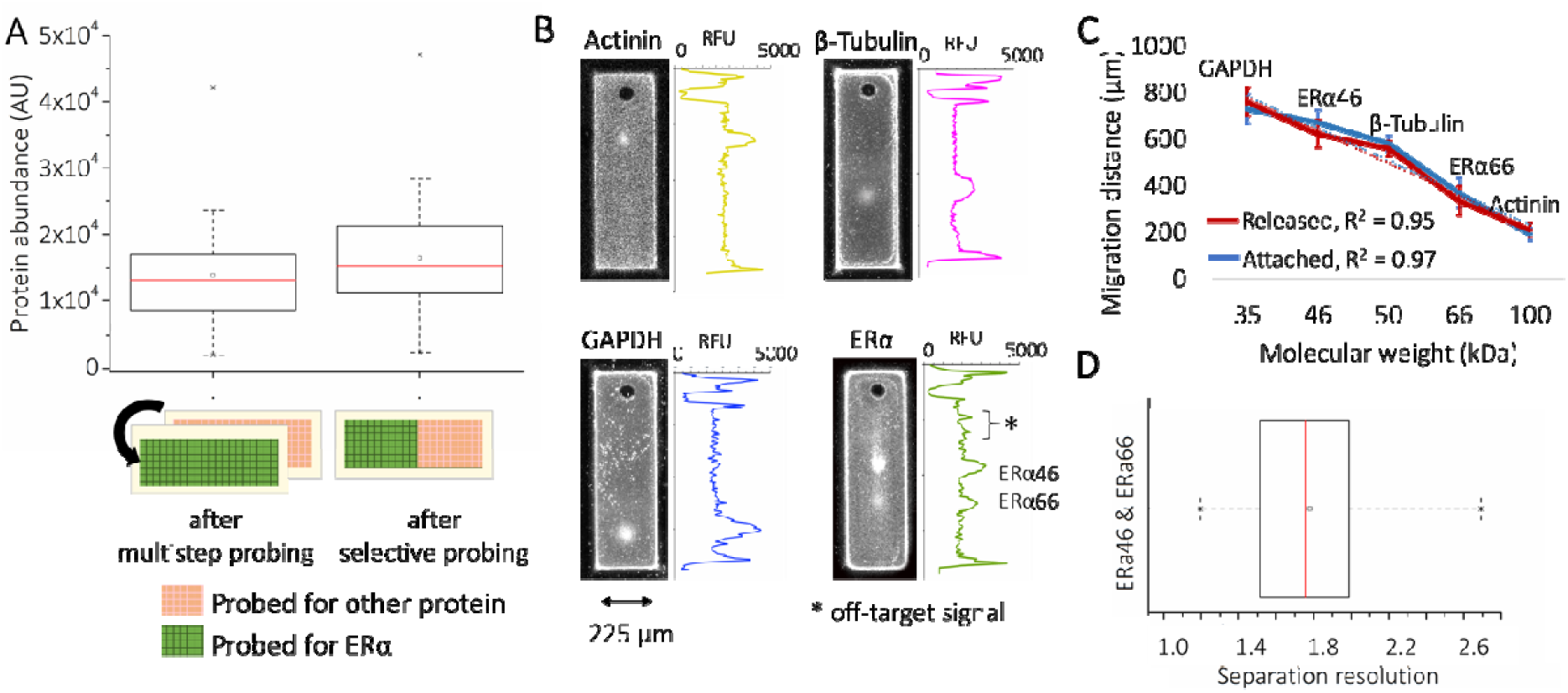
Characterization of ERα isoform expression in microparticle assay. **(A)** Measured ERα expression from the same array after a multistep probing (including one stripping round), and after probing of designated microparticles (p>0.05, n = 40 microparticles). **(B)** Fluorescence micrographs of microparticles for Actinin, β-Tubulin, GAPDH and ERα isoforms in MCF 7 cells. RFU, relative fluorescence units. The off-target peak (via ERα antibody) does not coincide with the ERα isoform bands. **(C)** Log-linear plot of species molecular weight against migration distance in 8%T PAG the fluorescently labeled species in panel A. (x-axis error bars within point size (± S.D., n = 3 separations); GAPDH, 35 kDa; ERα46, 46 kDa; β-Tubulin, 50 kDa; ERα66, 66 kDa; Actinin, 100 kDa). **(D)** Box plot demonstrating the separation resolution between ERα isoforms (n = 34 cells).

PAGE resolves two protein isoforms reactive to one ERα antibody probe: full-length (ERα66) estrogen receptor isoform (66 kDa) and truncated (ERα46) estrogen receptor isoform (46 kDa).[28,29] We validated the separation of ERα46 and ERα66 isoforms using three housekeeping proteins—Actinin (100 kDa), β-tubulin (50 kDa), and GAPDH (35 kDa)—as reference standards in a Ferguson plot analysis. We observed a linear relationship between migration distance and molecular mass for both the planar array and the suspended microparticles (R^2^ = 0.97, n = 121 microparticles and R^2^ = 0.95, n = 34 microparticles, respectively; Figure 2C). Separation resolution between the two ERα isoforms was 1.77 ± 0.33 (Figure 2D; n = 34 microparticles), which is considered to be baseline resolved and therefore quantitatively measurable.

### Dehydrated separations-encoded microparticles: Single-cell proteoform profiling

We next sought to explore unique performance gains possible in single-cell immunoblotting using the microparticle system. Specifically, we sought to understand analytical performance enhancements conferred by the isotropic shrinking of separation-encoded microparticles through dehydration of the microparticles from suspension. In a hydrogel microparticle that shrinks isotopically, we expect two potentially favorable scaling phenomena. First, we anticipate no sacrifice in separation resolution (SR, *SR* = *ΔL*/4*σ*), as both *ΔL* and *σ* should shrink by equal factors, assuming a detector with sufficient spatial resolution to resolve the final separation. At the hydrated state, the polymer network in separation-encoded microparticles is distributed uniformly. Shrinking develops upon evaporation of the aqueous phase (in this case water), initiating from the surface uniformly and resulting in a higher density polymer film formation at the outer interfaces.[30] Film formation may slow down the subsequent shrinking process under constant temperature and humidity levels, because the volume fraction of the polymer network in the film turns out to be much higher than in other portions of the gel.[30,31]. However, the water molecules inside the gel can still evaporate constantly even after the formation of the film layer since the molecules are small enough (Å-scale) compared to the pores in an 8%T 2.6%C polyacrylamide gel (nm-scale). After evaporation of all liquid content, the gel reaches a compact, uniform, and dehydrated state that does not impact *SR*.[32] Second, we expect improved analytical sensitivity of an immunoblot as the local concentration of fluorophores is inversely correlated with the volume (*L*^3^) of microparticles.[33]The average fluorescence signal increase (SI) can be calculated as follows, 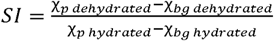, where χ is the intensity of the fluorescence signal, *p* is the location of the analyte peak, and *bg* is background.[34,35]

To understand the mechanism of hydrogel shrinkage, we performed single-cell immunoblots as described, then dehydrated the microparticle suspension by evaporation through heating on a hot plate. We measured a reduction of 83 ± 8 µm in microparticle length (950 µm to 866 µm; n = 250 microparticles) and 31 ± 5 µm in microparticle width (254 µm to 223 µm; n = 250 microparticles) suggesting isotropic shrinkage of each microparticle. The degree of circularity of the microwells was assessed (i.e.,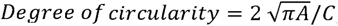, where *A* is the particle area including hole, *C* is the perimeter of the microwell.(36) Accordingly, a circular feature has a degree of circularity of 1.0, with non-circular features having values of <1.0. When comparing suspensions of hydrated microparticles to dehydrated microparticles, we observed no significant difference in the degree of circularity for the microwells (p-value <0.00001, n= 2881, 1823, 276, and 110, respectively; Figure S5B).

To next assess the impact of dehydration on the PAGE performance of separations-encoded microparticles, we assessed *SR* using GAPDH and β-tubulin in hydrated and dehydrated microparticles. We first considered the peak width of each target (*4σ*) probed and measured a 10% and 7% reduction in peak width (140 µm to 125 µm for GAPDH, 106 µm to 99 µm for β-tubulin; n = 121) (Figure 3A), consistent with observed shrinkage of the microparticle extents. We scrutinized any changes to ⍰*L* between the two markers stemming from dehydration-induced shrinkage of the separations-encoded microparticles.

**Figure 3.**
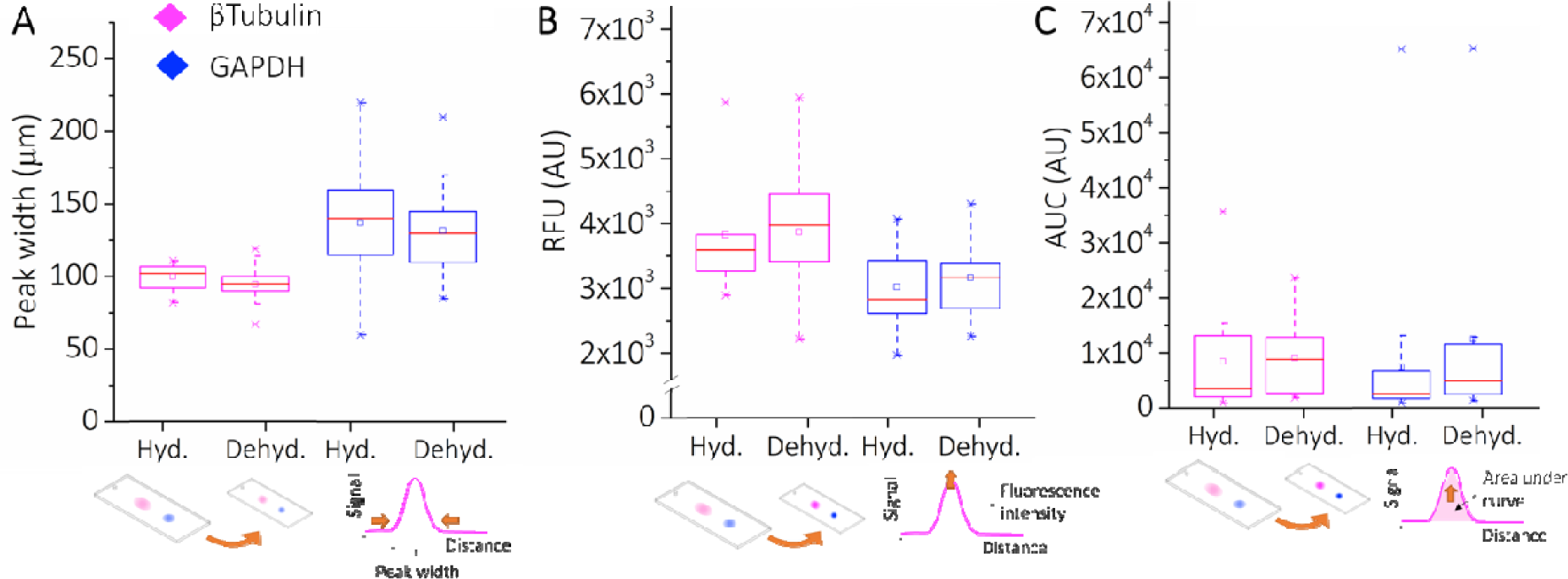
Characterization of microparticles with different morphologies. **(A)** Reduction in peak widths of β-tubulin and GAPDH were 10% and 7%, consistent with observed microparticle shrinkage. Mann-Whitney U-test p value was found to be lower than 0.05 (n = 121). **(B)** The median fluorescence signal intensity increase was 1.6x on the dehydrated microparticles compared to the hydrated microparticles (CV = 1.2, n = 121). **(C)** Quantified β-Tubulin and GAPDH expression in microparticles from U251 glioblastoma cells shows a median normalized AUC for dehydrated microparticles ∼1.3 to 1.7x higher than hydrated microparticles For all combinations, Mann-Whitney U-test p value was found to be lower than 0.05 (n = 121).

We found a median fluorescence signal intensity increase of 1.6x (CV = 1.2, n = 121) on the dehydrated microparticles relative to the hydrated ones (Figure 3B). According to the measured reduction in the dimensions of the dehydrated microparticles, fluorescence signal intensity increase (intensity/μm^3^) is expected to be 1.4x. Therefore, the measured increase is found to be in accordance with the calculated increase. Consequently, the measured *SR* values for the two markers were *SR* = 0.72 (CV = 22.86, n = 16 microparticles) in hydrated microparticle suspensions and *SR* = 0.72 (CV = 21.43, n = 62 microparticles) in dehydrated microparticles from suspension, which are not significantly different (two sample t-test p = 0.18), as anticipated from geometric arguments.

To understand the impact of microparticle shrinkage on analytical sensitivity and overall detection performance, we assessed the target signal (AUC) and the background signal. The impact of increased local concentration of fluorescence signal in dehydrated microparticles was characterized using GAPDH and β-tubulin. We found the median normalized AUC for dehydrated microparticles was ∼1.3 to 1.7x higher than hydrated microparticles (Mann-Whitney U-test p-value < 0.05 for each antibody type used, Figure 3C) (see SI). As described earlier, even modest improvements in the LOD facilitate higher coverage of the mammalian proteome (15% to 35%).

In understanding the effect of dehydration on the background signal, we compared the background fluorescence signal intensities obtained from hydrated/probed, hydrated/not probed, dehydrated/probed microparticles (Figure S5C). We probed separations-encoded microparticles with a secondary antibody (labeled with AlexaFluor 647) solution for 3 h and washed the excess solution for 1 h in TBST solution before imaging at 635 nm wavelength. Background fluorescence intensity of dehydrated/probed microparticles was ∼2x higher than hydrated/probed microparticles at the central regions, while this difference increases to be ∼40x relative to hydrated/not probed microparticles. The increase in the fluorescence intensity resulted in an enhanced SNR, defined as the ratio of the average fluorescence signal minus the mean background signal to the standard deviation of the background.(37) The noise on the dehydrated microparticles was noted to decrease by 1.5x (median SNR = 9.8, CV = 0.8, n = 392 microparticles) relative to the hydrated microparticles, SNR = 6.4 (CV = 0.4, n = 342 microparticles). The lower noise in dehydrated microparticles yields an improved analytical sensitivity. This result is in accordance with our observations regarding the effect of microparticle shrinkage on the target detection signal. The geometry-induced performance changes made target signal detectable in 17% more dehydrated microparticles versus detectable in hydrated microparticles. Reduced noise in the separations-encoded microparticles is attributed to the enhanced mass-transport of antibodies during the washout step, although dehydration (shrinkage) process increases the signal intensity in microparticles.

Next, we measured ERα protein isoform expression differences using the separations-encoded microparticles, ERα46 (46 kDa) and ERα66 (66 kDa) isoforms, in MCF 7 (estrogen sensitive), MDA MB 231 (estrogen resistant), and HEK 293 (non-expressing) cells at different confluency levels. For the estrogen-sensitive cell line, we expect an inverse correlation between ERα46 expression and ERα66 expression if hormonal resistance increases with ERα46 expression.[28] We first confirmed that the gradual increase in cell confluency levels agrees with the gradual increase in housekeeping protein expression levels for all cell lines (Figure 4A and Figure S6A). Figure 4A shows no increase in housekeeping protein levels, which we attribute to normalizing signal by the number of cells assayed per day. On the basis of separations-encoded microparticle assay results, we compared SNR of ERα46 and ERα66 in MCF 7 cells and found that the average of both SNR values was above SNR = 3 threshold. Particularly, the SNR average was 4.79 for ERα46 (n = 385) and 6.41 for ERα66 (n = 93) (Figure 4B). Importantly, AUC analysis in separations-encoded microparticles revealed a 2.8x increase in truncated isoform (ERα46) and 6.4x decrease in full-length isoform (ERα66) over a 14-day period in estrogen sensitive cells (Figure 4C, n = 478 cells); therefore, confirmed that higher confluency increases ERα46 expression that suppresses the expression of ERα66 in MCF 7 cells.[28,29] Surprisingly, separations-encoded microparticles reported minute levels of ERα46 isoform-expressing cells in the estrogen resistant cell line, while this population remained masked in slab gel westerns (Figure S7). ERα66 isoform was not detected in estrogen resistant cell line as it is in the class of highly invasive phenotype that reportedly lacks ERα66 isoform.[29] Neither of the isoforms was detected in the non-expressing cell line, which has been used as a negative control in this experiment.

**Figure 4.**
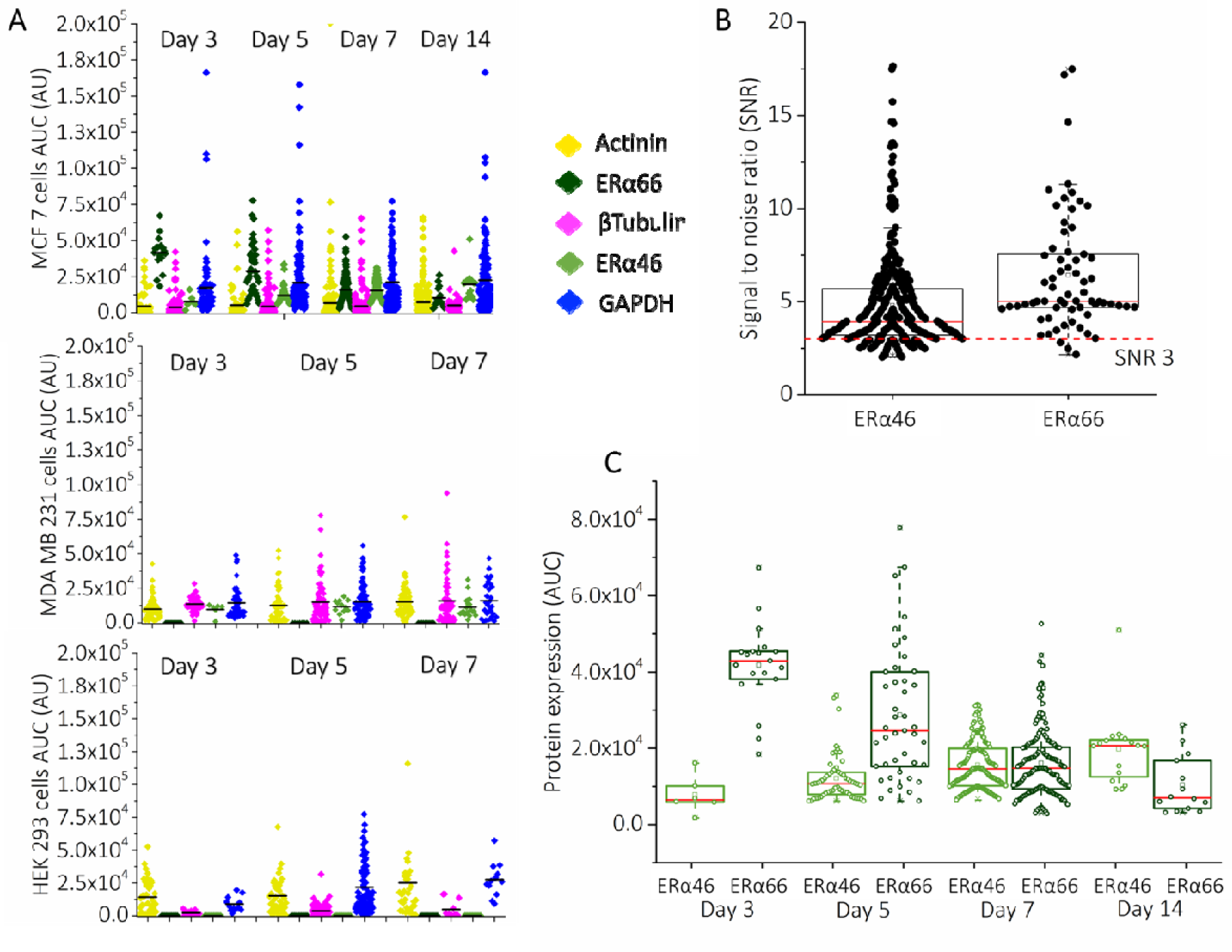
Expression of ERα isoforms change over confluency of cell culture. MCF 7 cells are shown as the model ERα positive organisms, whereas MDA MB 231 and HEK 293 serve as the ERα negative control cell lines. **(A)** Color-coded beeswarm graphs show single-cell protein measurements in subsequent cell culturing days for MCF 7, MDA MB 231, and HEK 293 cell lines. The black bars present the median protein expression level for each group. **(B)** Signal-to-noise ratio (SNR) for ERα isoforms. Red dashed line presents SNR = 3, above which protein quantification was employed for all measurements, n_total_ = 447. **(C)** The grouped box plots show fluctuations in ERα isoform expression over 14 days in MCF 7 cells (n = 478 cells). Microparticles reported an gradual increase in the number of cells expressing ERα46 (green), while the expression of ERα66 (blue) dropped gradually over 14 days.

Quantification of ERα expression changes in MCF 7 cells benefited from the use of releasable separations-encoded microparticles in two key aspects. First, enhanced mass transport in microparticles helped to reduce the total immunoprobing time from ∼50 hours to ∼14 hours, even though we employed two probing and stripping rounds for a total of 5 protein species. Second, we achieved to discriminate minor cell populations representing ERα46 expressing MCF 7 cells, (e.g. 6 cells on day 3, 54 cells on day 5, and 163 cells on day 7 from a large cell population of 478 cells, see SI).

## Experimental

### Chemicals and reagents

Acrylamide/bis-acrylamide, 30% (wt/wt) solution (Sigma-Aldrich, cat. no. A3699), sodium dodecyl sulfate (SDS) (cat. no. L3771), sodium deoxycholate (NaDOC) (cat. no. D6750), Triton X-100, N,N,N⍰, N⍰-tetramethylethylenediamine (TEMED) (cat. no. T9281), ammonium persulfate (APS) (cat. no. A3678), and bovine serum albumin (BSA) (cat. no. L3771) were purchased from Sigma Aldrich. Goat anti-GAPDH primary antibody (Sigma-Aldrich, cat. no. SAB2500450), mouse anti-βTubulin (Genetex, cat no. GTX11312), rabbit anti-ERα SP1 (Thermo Scientific, cat. no. RM-9101-S0), rabbit anti-actinin (Cell Signaling, cat no. 6487) were purchased from the suppliers. N-[3-[(3-Benzoylphenyl)-formamido]propyl] methacrylamide (BPMAC) was synthesized by PharmAgra Laboratories. Tris-glycine (10×) EP buffer was procured from Bio-Rad (25 mM Tris, pH 8.3; 192 mM glycine, cat. no. 1610734). Phosphate-buffered saline was acquired from VWR (10× PBS) (cat. no. 45001-130). Petroleum jelly was purchased from Cumberland Swan Petroleum Jelly (cat. no. 18-999-1829). Tris-buffered saline with Tween (TBST) was obtained from Santa Cruz Biotechnology (20× TBST) (cat. no. 281695). Deionized water (18.2 MΩ) was obtained from an Ultrapure Millipore filtration system.

### Fabrication of SU-8 mold and separations-encoded microparticles

The SU-8 mold fabrication was performed by following the manufacturer’s instructions (MicroChem). Approximately 7000 microparticles (250×1000×40 µm) can be fabricated from a 70×48-mm mini-slab polyacrylamide gel. Each microparticle contained a microwell (30 µm in diameter), a separation lane (950 µm in length) and 50-µm wide perforations defining the microparticles. 8%T, 2.6%C polyacrylamide gel containing 5 mM BPMAC was layered on an acrylate-silanized microscope glass slide by the help of the SU-8 mold. The gel was synthesized by chemical polymerization using 0.08% APS as the initiator and 0.08% TEMED as the catalyst. The gels were incubated in distilled water for 10 min at room temperature prior to releasing the glass slide from the SU-8 mold. After following single-cell western blotting procedure, microparticles were released from the glass slide by shearing using a razor blade when the microparticles are in the hydrated state. Thanks to the perforations, individual particles can be released easily.

### Cell culture

Breast adenocarcinoma (MCF 7), invasive breast adenocarcinoma (MDA MB 291), and embryonic kidney (HEK 293) cell lines were purchased from ATCC. The MCF 7 cells were cultured in RPMI 1640 media (ThermoFisher Scientific, cat. no. 11875093) supplemented with 10% Charcoal-stripped serum (Sigma-Aldrich, cat. no. F6765), and 1% penicillin/streptomycin (ThermoFisher Scientific, cat. no. 15140122). MDA MB 231 cells were cultured in the same culture media with MCF 7 cells. U251-GFP cells were cultured in high glucose DMEM (Life Technologies, cat. no. 11965) supplemented with 1% penicillin/streptomycin, 1× MEM nonessential amino acids (11140050), 1 mM sodium pyruvate (Life Technologies, cat. no. 11360-070), and 10% FBS. and HEK 293 cells were cultured in the same culture media with U251-GFP cells, except 10% Charcoal-stripped serum (Sigma-Aldrich, cat. no. F6765) was used instead of 10% FBS. All cells were grown in a humidified atmosphere containing 5% CO_2_ at 37 °C. All cells were cultured in flasks.

### Single-cell western blotting using separations-encoded microparticles

#### Single-cell resolution western blotting

Approximately 1×10^6^ U251-GFP cells were settled by gravity into the wells located on microparticles for 10 min, and the gel was washed three times with 1 mL 1× PBS to remove excess cells off the gel surface. Then, settled cells were lysed by directly pouring the RIPA-like lysis buffer at 55 °C over the slide. During cell lysis, portions of each single-cell lysate are diluted and lost by diffusion and convection, as has been characterized in detail by our group.[16] We optimized the lysis and electrophoresis times to achieve a separation of <30% mass difference between two neighboring proteins in the assay. Cell lysates were electrophoresed at 40 V cm^−1^ for 30 s, and immediately after protein bands were immobilized by UV activation (Lightningcure LC5, Hamamatsu) of the BPMAC. The microparticle array was incubated in TBST buffer overnight at room temperature.

#### Immunoprobing and imaging of separation-encoded microparticles after single-cell western blotting

Donkey Anti-Goat IgG (H+L) Cross-Adsorbed Cross-Adsorbed Antibody, Alexa Fluor 555-labeled antibody (cat. no. A21432), Donkey Anti-Rabbit IgG (H+L) Cross-Adsorbed antibody, Alexa Fluor 555-labeled (cat. no.A21432), and Donkey Anti-Rabbit IgG (H+L) Cross-Adsorbed Secondary antibody, Alexa Fluor 647-labeled (cat. no. A31573) were purchased from Thermo Fisher Scientific. After the electrophoretic separation and UV-induced covalent attachment of proteins to benzophenone in microparticles, the gel was washed for overnight in 1× TBST on an orbital shaker. Separations-encoded microparticles were incubated in 30 μL of a 1:10 dilution of primary antibodies in 1× TBST with 2% BSA solution for three hours, and washed three times for 20 min in 1× TBST. Secondary antibodies were incubated for two hours in a 1:10 dilution and gels were washed three times for 20 min in 1× TBST and dried using nitrogen stream. After the immunoprobing, the hydrated microparticle array was released from the glass slide surface using a razor blade. During this process, microparticles were completely peeled off from the glass surface without any remaining parts attached to the surface. Separations-encoded microparticles can be imaged in the released format or in the array format. For all cases (attached, released, hydrated, and dehydrated states), we imaged the gel particles using a fluorescence microarray scanner (Molecular Devices, Genepix 4300A) with an Alexa-Fluor 555 filter (532 laser excitation, 650 PMT), and an Alexa-Fluor 647 filter (647 laser excitation, 550 PMT).

#### Data analysis and simulations

Protein bands were quantified using in-house developed MATLAB scripts as described in Kang et al.[16] Peak widths were characterized by Gaussian curve fitting in MATLAB (R2017b, Curve Fitting Toolbox). The integrated intensity of a marker was calculated if its R^2^ value was larger than 0.7 for the given region of interest. For all quantified results, the protein peaks with Gaussian fitting R^2^ value larger than 0.7 and SNR value larger than 3 were analyzed. Electric field distribution simulations were performed in COMSOL Multiphysics 5.3a (Burlington, MA) to characterize protein mobility during electromigration in separations-encoded microparticles. A 2D asymmetric model was used. The gel width was 250 µm and the length was 1000 µm. The diameter of the microwell was set to 30 µm. The maximum and minimum mesh element sizes were 30 and 0.3 µm, respectively. The model was solved in stationary mode by applying a constant electric field through the matrix, where hydrogel and solution conductivities were set to 4304.0 μS m^−1^ estimated experimentally.[38] The goal of the simulations was to estimate the electric field instabilities caused by discontinuous surface area created by arrayed microparticles.

### Slab gel western blotting

#### Imaging of living cells

The confluency of the cells was determined by imaging the culture dishes in bright field using an Olympus IX71 inverted fluorescence microscope equipped with an EMCCD camera (iXon3 885, Andor, Belfast, Ireland), motorized stage (Applied Scientific Instrumentation, Eugene, OR), and automated filter cube turret controlled through MetaMorph software (Molecular Devices, Sunnyvale, CA).

#### Cell lysate preparation for slab gel western blotting

At the desired confluency level, cells were trypsinized using 1 mL of trypsin EDTA solution for 1-2 min at 37 °C and 4 mL culture medium was added to the 25 cm^2^ (T-25) culture flask. After centrifuging the cells at 1000 rpm for 5 min in a 15 m tube, the culture medium was aspirated. Cells were resuspended in 1 mL PBS and counted using trypan blue stain. Cells were washed two times more with ice-cold PBS to remove all the loosely bound serum proteins in the media before adding 1mL of ice-cold protease inhibitor-contained RIPA buffer (0.15 mmol L^−1^ NaCl, 5 mmol L^−1^ EDTA, 1% Triton-X 100, 10 mmol L^−1^ Tris-HCl. Cells were incubated on ice for 15 min while vortexing in every 5 min. The suspension was transferred to an Eppendorf tube and centrifuged at 25,000 × g for 30 min at 4 °C. The supernatant containing 30 µg mL^−1^ of cellular proteins were used in slab gel western blotting.

#### Slab gel western blotting

4-12% gradient gel (Novex WedgeWell Tris-Glycine, cat. no. XP04125BOX) and a Bio-Rad electrophoresis chamber connected to a Bio-Rad high voltage power supply set at 200 V were utilized for protein separation from cell lysates. Each well was loaded with 30 µg protein labeled with a 9:1 mixture of 4× Lamelli buffer and 10× NuPAGE reducing agent at 1:1 ratio. After the electrophoresis, Pierce PowerBlot Rapid Transfer System (ThermoFisher Scientific, cat. no. PB0112) was used to transfer the proteins to a PVDF membrane according to the manufacturer’s instructions. The PVDF membrane was blocked in 5% BSA (w/v) solution for 30 min and was incubated in 10 mL solution with a 1:1000 dilution of primary antibodies in 1× TBST with 5% BSA solution for overnight at 4 °C. The membrane was rinsed three times with TBST for 10 min each round. Secondary antibodies were incubated in a 5% BSA solution with a 1:10000 dilution for 1 h and PVDF membrane was rinsed using TBST three times with TBST for 10 min each round. After the rinsing step, Western Lightning (PerkinElmer, cat. no. NEL120E001EA) was utilized as described in manufacturer’s protocol for obtaining chemiluminescence images using a Chemidoc XRS system (Bio-Rad, cat. no. 170-8265).

## Conclusions

We have designed, developed, and applied separations-encoded microparticles to advance performance and utility of single-cell immunoblotting, which has been limited by microscale mass transport considerations particularly during immunoprobing. Mass transport limitations are a bottleneck of immunoassays, requiring substantial dedication of assay time to the probing and washout steps. Separations-encoded microparticles reduce mass transport restrictions, boost analytical performance, and bring handling flexibility by combining normally disparate microarray and microparticle formats. Furthermore, protein immunoblotting with separations-encoded microparticles confers selectivity suitable for isoform detection while allowing selective probing and multiplexing by isolation individual cell readouts. Merging microparticles with biomolecular separations surmounts measurement challenges where (1) immunoassays are insufficient owing to limited probe selectivity and (2) powerful mass spectrometry tools lack analytical sensitivity and throughput. The unique capabilities of the separations-encoded microparticles should provide a tunable, versatile format for cytometry.

## Acknowledgements

We thank all the Herr lab members for helpful discussions and suggestions throughout the study. We thank Dr. John J. Kim for fruitful discussions on ERα isoform separations and selection of regarding cell lines. This work was supported by the grant from the United States National Institutes of Health (NIH Grant R21EB019880 to A.E.H.) and Chan Zuckerberg Biohub.

## Supporting Information

### Quantitative microparticle analysis of ER protein isoforms from single cells

Estrogen receptor, ER, is the main contributor to the development of hormone resistance against the therapy in *circa* 70% of breast tumors.(29) Particularly, truncated (ERα46) estrogen receptor isoform is reported to enhance hormone therapy responses by inhibiting the expression of full-length (ERα66) estrogen receptor isoform.(28) The relation between cell confluency and ERα expression in hormone-sensitive cell lines has remained unclear, although this finding may suggest new strategies in therapies. Elucidating mechanisms of confluency-dependent expression of these isoforms requires quantitative measurement of ERα isoforms. We hypothesized that cell confluency level would impact expression levels of ERα46 protein in estrogen-sensitive cells, while no expression would be detected in estrogen resistant and non-expressing cell lines. We used separations-encoded microparticles to address the confluency-dependence of ERα46 and ERα66 expression levels in estrogen sensitive (MCF 7) over 14 days. We performed slab gel Western blotting assay to validate the microparticle assay results (Figure S5B and Figure S5C). A similar trend in ERα isoform expression levels was observed in slab gel Western analysis, except that no expression could be detected on day 3 of MCF 7 cells unlike microparticle analysis results. We did not observe any detectable signal from ERα isoforms expressed in MDA MB 231 in slab gel Westerns, although microparticles reported 6, 12, 20 cells expressing ERα46 on days 3, 5, and 7, respectively (Figure S7) (n_total_ = 38). We did not observe any ERα isoform signal in HEK 293 cells as shown in Figure S7. Slab gel Western blot analysis confirmed our major finding on ERα expression level changes in MCF 7 cells using microparticle assay, which provided more insight into isoform expression in cell subpopulations.

### Characterization of the electric field distribution in microparticle assay

A uniform electric field distribution is compulsory for maintaining repeatable and comparable protein separation in microparticle arrays. In a typical workflow, we apply an electric field across the microparticle array to draw proteins through the microwell wall into the central region of microparticles. The uniformity of the field can be ensured by immersing the unidirectionally placed microparticles on top of a silanized glass plate in a conductive buffer where two parallel platinum electrodes are placed in to apply the electric field. We characterized the uniformity of electric field using two approaches: (1) we simulated electric field distribution in the assay, (2) we tracked the peak position displacement of β-tubulin (50kDa) and GAPDH (35kDa) proteins as a function of time in microparticles placed on a glass slide. Firstly, the electric potential in the microparticle assay was found to be uniform and unaltered between the electrodes according to numerical simulations (Figure S8). This was an expected outcome because polyacrylamide gel must not result in a discontinued conductivity along the separation axis as long as soaked in the buffer solution during electrophoresis.(38) Simulation result was also confirmed by electrophoresis of β-tubulin and GAPDH proteins obtained from U251 cells (Figure S4). We also tested electrophoretic mobility of two proteins (β-tubulin and GAPDH) on non-silanized glass slides, as it is known that silanization process does not always result in a uniform monolayer of silane groups on the glass slide.(16) Comparison of measured mobilities shown in Figure S4 suggest that silanization state does not have a statistically significant impact on the electrophoretic mobility of the proteins (two sample t-test, p = 0.0001, n_silanized glass_ = 20, n_non-silanized glass_ = 20).

### Cell-occupancy dependent fluorescence signal changes in separations-encoded microparticles

We compared the antibody fluorescence intensities in multiple-cell occupancy microwells and single-cell occupancy microwells at the central region of microparticles. The mean antibody fluorescence intensity from two-cell occupancy microparticles was ∼5.5 times higher than that of single-cell occupancy microparticles (CV = 0.28, n = 10 microwells). The intensity was not doubled (therefore not linear) in two-cell occupancy microparticles due to the cell-size related bias. This result is in accordance with our previous findings.(16)

### Characterization of the fluorescent signal intensity distribution

As we discussed in the manuscript, shrinking effect led to increase of the fluorescence signal as the equal number of fluorescence molecules were packed in a smaller volume after dehydration. Interestingly, we observed significantly higher signal at the edges of the central region of separations-encoded microparticles. The edge effect was observed 2.35x higher in hydrated/not probed particles, 5.35x higher in dehydrated particles, 8.11x higher in hydrated particles (Figure S5C). We attribute this occurrence to hydrogel pore size formation at the outer boundaries during polymerization. Polymerization reaction rate is slower at the edges because of the higher oxygen concentrations (oxygen diffuses in the particles passing through the edges first).(39) Reduced reaction rate leads to formation of larger pores, in which fluorescent molecule concentration can be higher.(40) The edge effect did not interfere with the of protein measurements since the background intensity in the central region of the particles was uniformly distributed.

---

**Figure S1.**
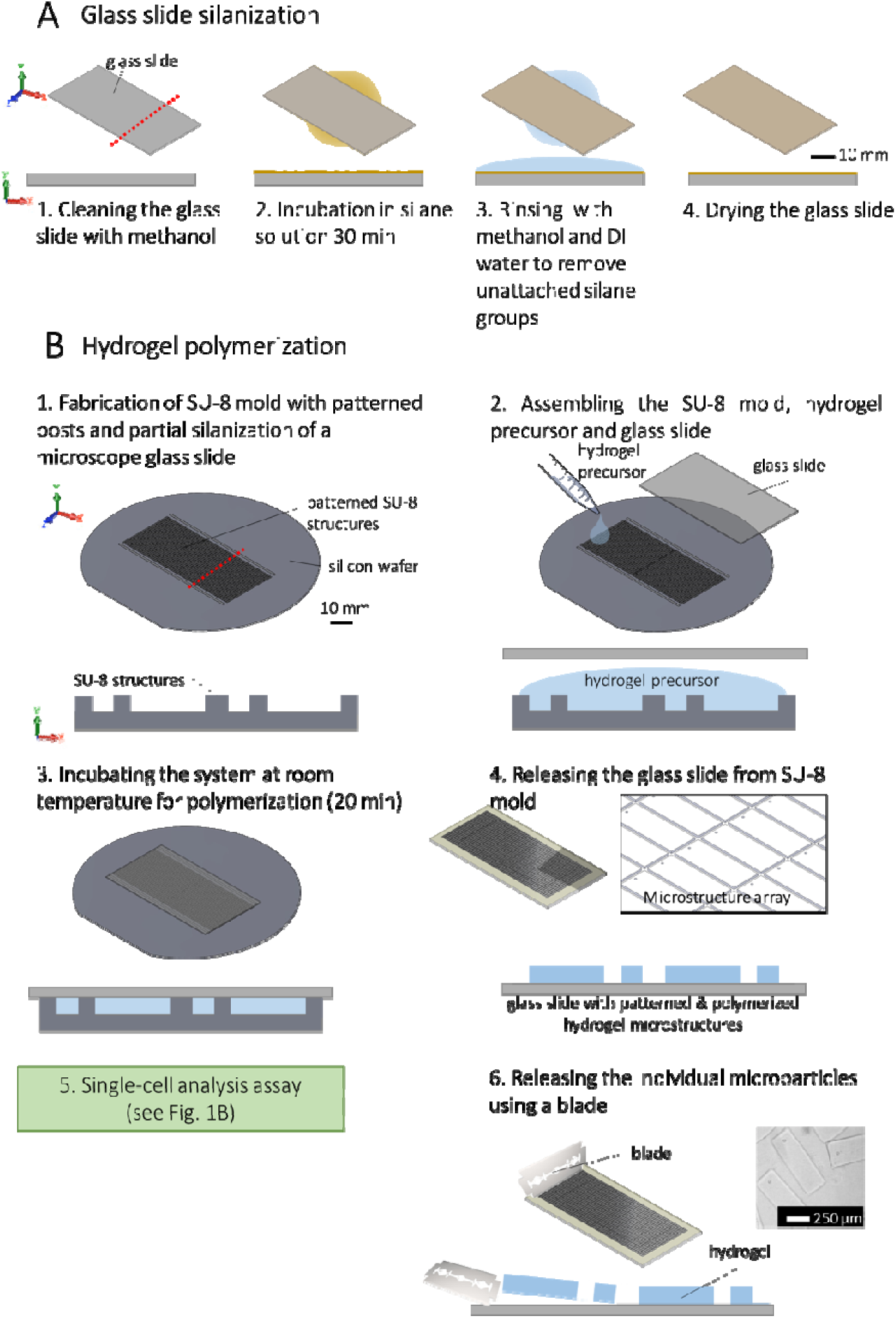
Conceptual description of the workflow of separations-encoded microparticle fabrication. **(A)** A microscope slide is silanized to attach microparticle array on the glass. **(B)** Silanization process is followed by polyacrylamide gel synthesis based on chemical polymerization. Hydrogel precursor is sandwiched between the silanized microscope slide and a silicon wafer with SU-8 posts. After releasing the microscope slide with microparticle patterns from the silicon wafer, single-cell analysis is employed. Individual microparticles can be released from the microscope slide using a razor blade.

**Figure S2.**
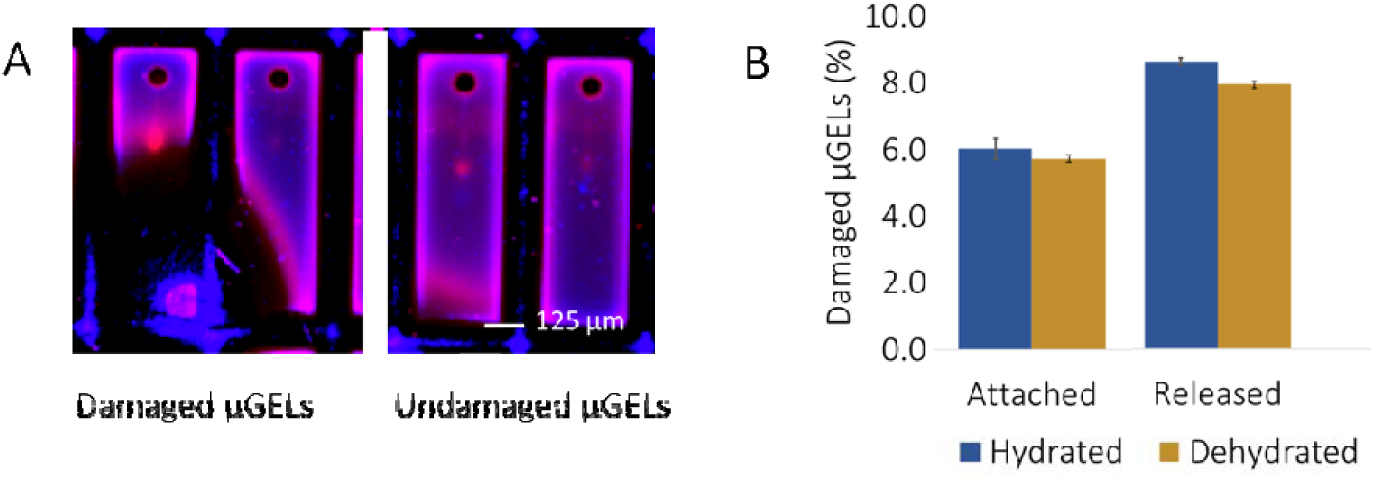
Damage analysis in microparticles. **(A)** False-color fluorescence image of damaged and undamaged microparticles. **(B)** Bar graph shows the percentage of damaged microparticles at hydrated and dehydrated states after and before releasing, p < 0.0001 (two sample t-test), n = 2483 microparticles.

**Figure S3.**
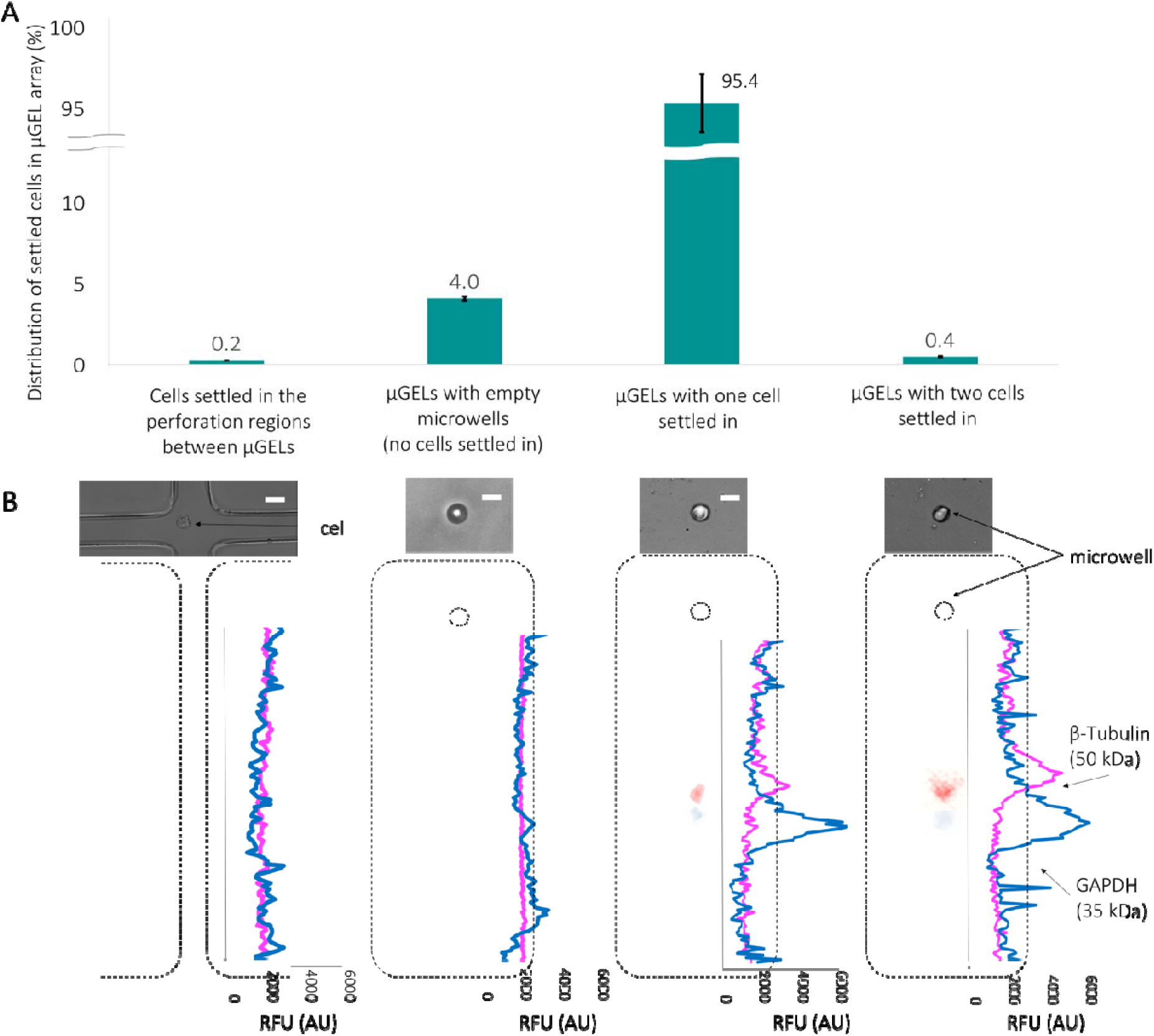
Characterization of cell settling in the microparticle array. **(A)** Bar graph summarizes the distribution of settled cells in microwells (n = 3 devices with 3500 units each). **(B)** Bright field images (top) and fluorescence intensity graphs (bottom) of microparticles. Separation of β-Tubulin and GAPDH proteins are shown and the mean antibody fluorescence intensity from two-cell occupancy microparticles was ∼5.5x higher than that of single-cell occupancy microparticles (CV = 0.28, n = 10 wells). The intensity was not doubled (not linear) in two-cell occupancy microparticles due to the cell-size related bias.

**Figure S4.**
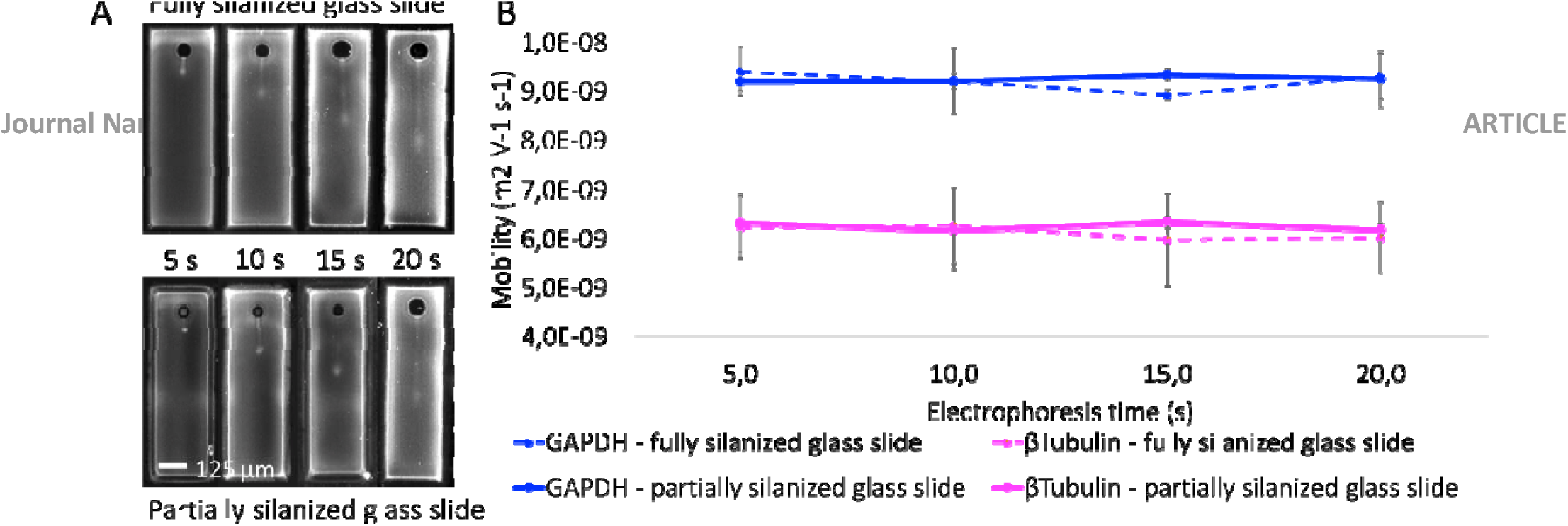
Mobility measurements for β-Tubulin and GAPDH proteins are performed in 8%T 2.6%C polyacrylamide gel microparticles fabricated on partially and fully silanized glass slides. **(A)** Montage of fluorescence micrographs of β-Tubulin at different electrophoresis times on partially and fully silanized glass slides. **(B)** Changes in β-Tubulin (50 kDa) and GAPDH (35 kDa) mobilities in microparticles attached on partially and fully silanized glass slides, n_total_ = 40 microparticles.

**Figure S5.**
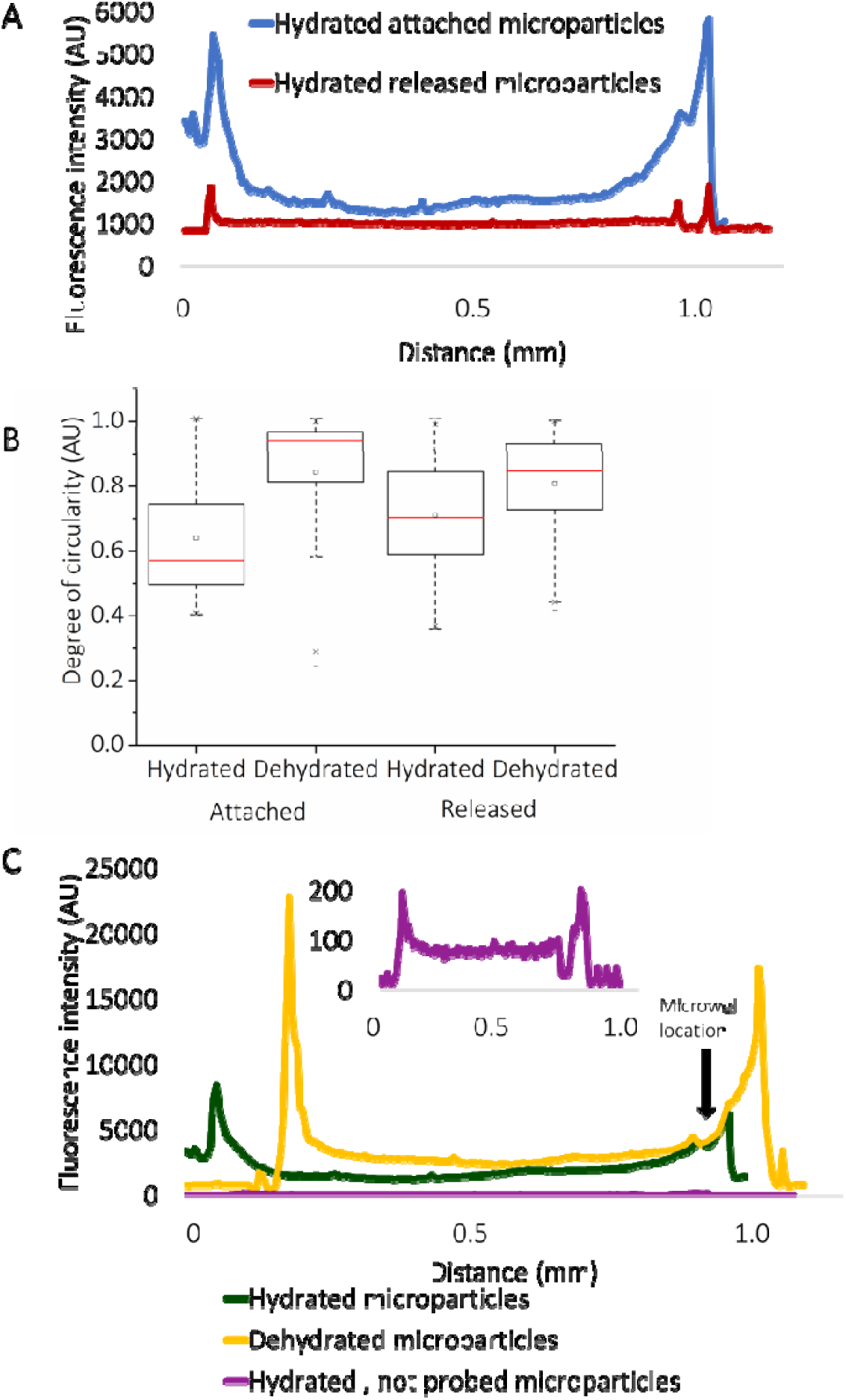
Comparison of fluorescence intensities in microparticles **(A)** Background intensity reduction in released microparticles suggests the geometry-enhanced mass transport during antibody probing and washout steps. **(B)** Characterization of degree of microwell circularity changes in combinations of hydrated, dehydrated, attached, and released states. Surface area normalized to attached hydrated condition is shown in the box plot, p < 0.00001 between all groups, n_total_ = 5090. **(C)** Fluorescence intensity graph shows signal intensity changes in hydrated/probed, hydrated/not probed, dehydrated/probed microparticles The inset shows a close-up intensity profile of hydrated/not probed microparticles.

**Figure S6.**
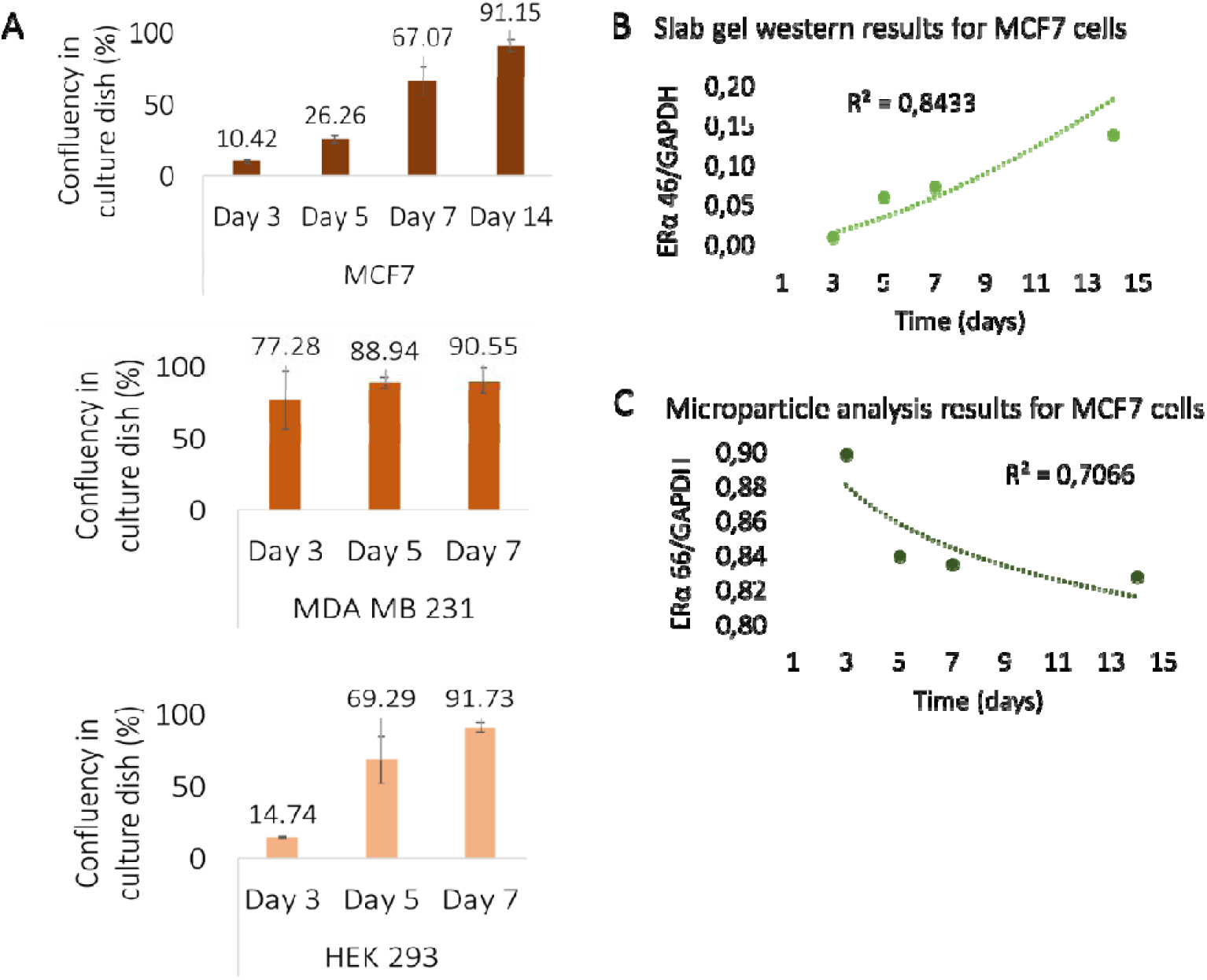
Cell confluency level changes in three cell lines and change of ERα isoform expression over confluency of MCF 7 cells in slab gel westerns. **(A)** The bar graph shows the confluency levels of estrogen sensitive (MCF 7) and resistant (MDA MB 231) breast cancer cell lines as well as HEK 293 cells over 7 and 14 days. MDA MB 231 and HEK 293 cells reached > 90% confluency after 7 days of cell culture. Slab gel western blots (15 mg protein) were performed to confirm microparticle assay results. The scatter graph of **(B)** ERα46 and **(C)** ERα66 to GAPDH expression ratio shows measured increase and decrease in protein expression levels over 14 days (n = 2 slab gel western runs).

**Figure S7.**
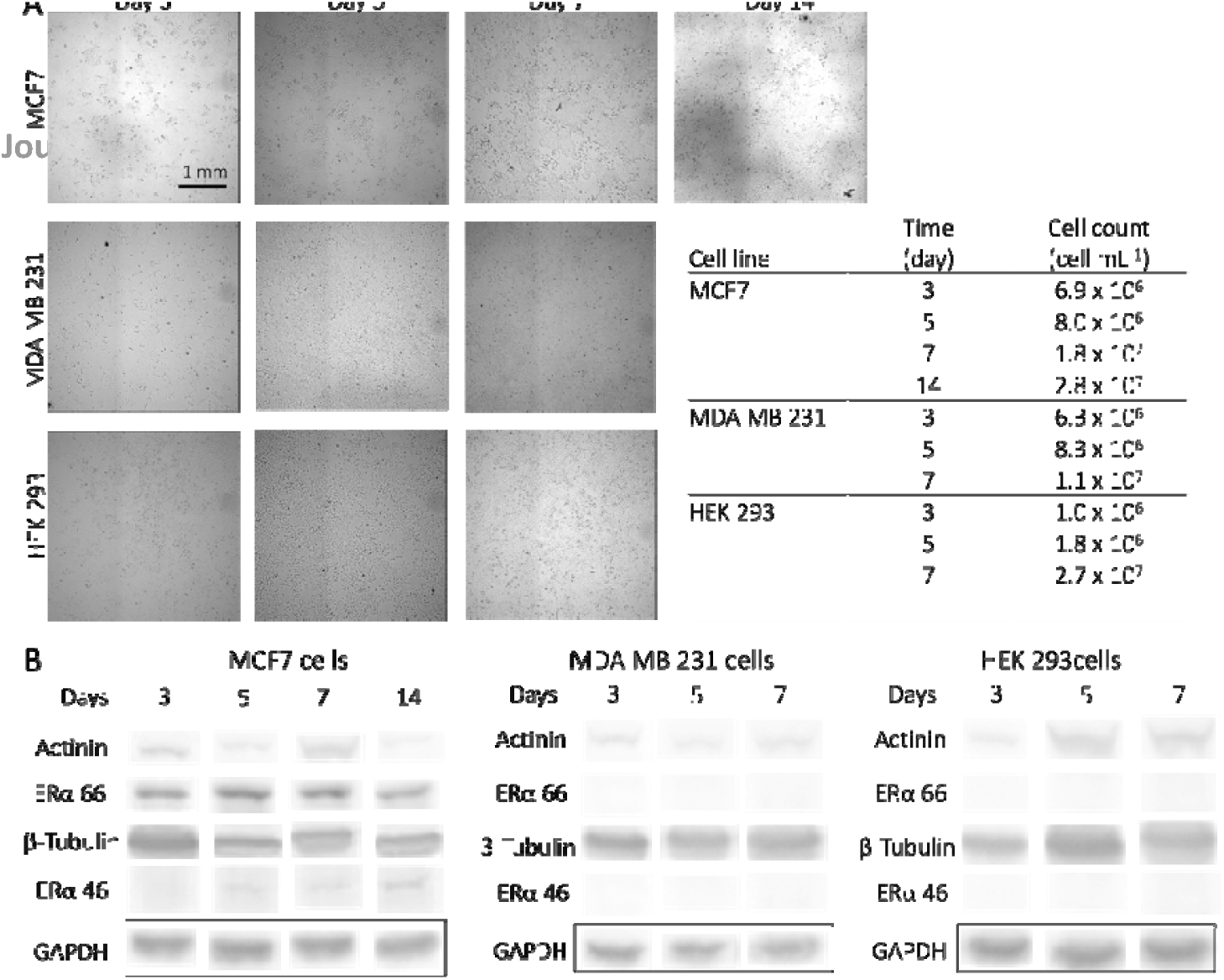
Comparison of cell confluency and ERα expression levels. **(A)** Bright-field images of MCF 7, MDA MB 231, and HEK 293 cells in corresponding culture days and counted cells per mL that are used to calculate confluency levels. **(B)** Identification of ERα isoforms in MCF 7, MDA MB 231, and HEK 293 cells by conventional slab gel Western blotting. Western blotting images were used in quantification of ERα isoforms in MCF 7 cells. No detectable signal was observed in MDA MB 231, and HEK 293 cells, as expected.

**Figure S8.**
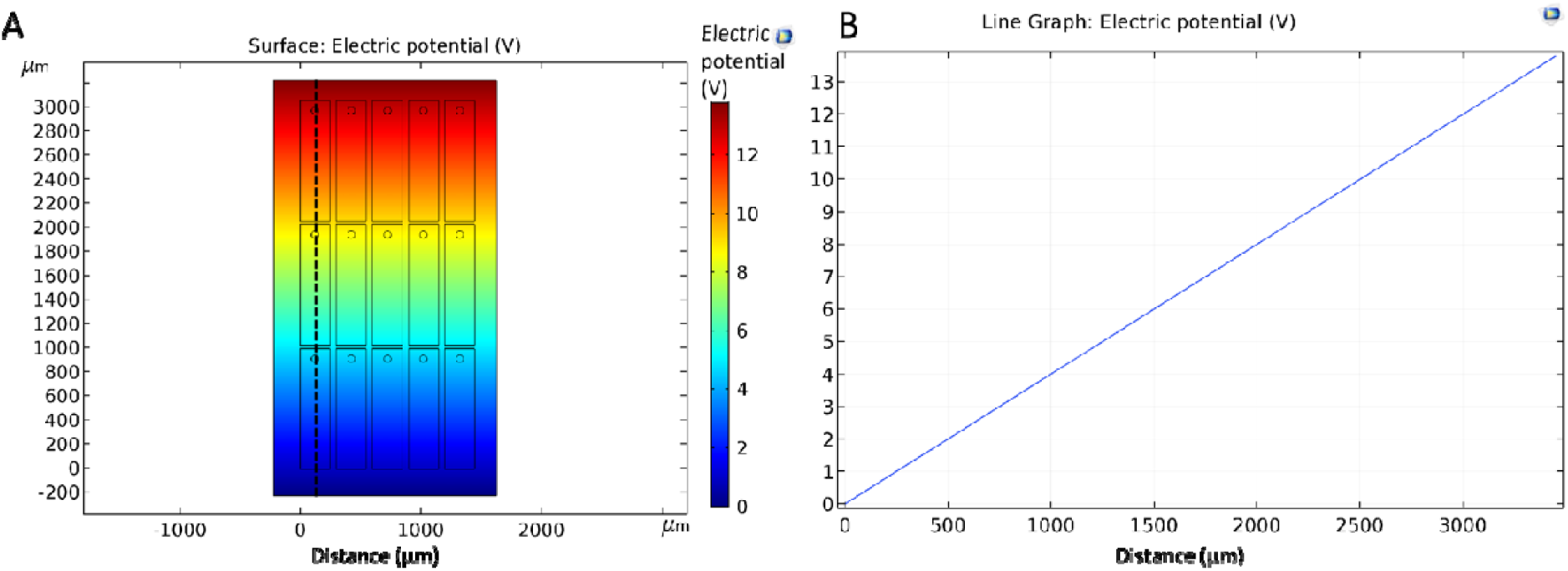
Finite element simulation of electric field distribution. Electric currents module of COMSOL Multiphysics software was operated by applying steady-state constant electric field through the matrix, where hydrogel and solution conductivities were set to 4304.0 μS m^−1^ estimated experimentally. **(A)** A section of microparticle array geometry and the cross section along which the electric field was simulated. **(B)** Electric field distribution in the section at dashed black line in panel A shows that microparticles do not disturb electric field distribution.

## References

1 C. Schubert, Nature 2011, 480, 133.

2 P.K. Chattopadhyay, T.M. Gierahn, M. Roederer, J.C. Love, Nat. Immunol. 2014, 15, 128.

3 W.K. Smits, O.P. Kuipers, J.W. Veening, Nat. Rev. Microbiol. 2006, 4, 259.

4 L. Luo, X. Li, R.M. Crooks, Anal. Chem. 2014, 86, 12390–12397.

5 PS. Phadtare, V.P. Vinod, P.P Wadgaonkar, M. Rao, M. Sastry, Langmuir 2004, 20, 3717–3723.

6 P. Mao, R. Gomez-Sjoberg, D. Wang, Anal. Chem. 2012, 85, 816–819.

7 M. Sevecka, G. MacBeath, Nature Methods 2006, 3, 825–831.

8 A.H. Ng, M.D. Chamberlain, H. Situ, V. Lee, A.R. Wheeler, Nature Commun. 2015, 6, 7513.

9 T.M. Ward, E. Iorns, X. Liu, N. Hoe, P. Kim, S. Singh, S. Dean, A.M. Jegg, M. Gallas, C. Rodriguez, M. Lippman, Oncogene. 2013, 32, 2463.

10 C. Lombard-Banek, S.A. Moody, P. Nemes, Angew. Chem. Int. Ed. 2016, 55, 2454–2458.

11 S.C. Bendall, E.F. Simonds, P. Qiu, D.A. El-ad, P.O. Krutzik, R. Finck, R.V. Bruggner, R. Melamed, A. Trejo, O.I. Ornatsky, R.S. Balderas, Science 2011, 332, 687–696.

12 M.H. Spitzer, G.P. Nolan, Cell, 2016, 165, 780–791.

13 T.A. Duncombe, A.E. Herr, Lab Chip 2013, 13, 2115–2123.

14 B. Chen, S. Lim, A. Kannan, S.C. Alford, F. Sunden, D. Herschlag, I.K. Dimov, T.M. Baer, J.R. Cochran, Nat. Chem. Biol. 2016, 12, 76.

15 C. Zhang, H.L. Tu, G. Jia, T. Mukhtar, V. Taylor, A. Rzhetsky, S. Tay, Sci. Adv. 2019, 5, eaav7959.

16 C.C. Kang, K.A. Yamauchi, J.V. Vlassakis, E. Sinkala, T.A. Duncombe, A.E. Herr, Nature Protoc. 2016, 11, 1508.

17 R. Fan, O. Vermesh, A. Srivastava, B.K. Yen, L. Qin, H. Ahmad, G.A. Kwong, C.C. Liu, J. Gould, L. Hood, J.R. Heath, Nature Biotechnol. 2008, 26, 1373.

18 A.J. Hughes, D.P. Spelke, Z. Xu, C.C. Kang, D.V. Schaffer, A.E. Herr, Nature Met. 2014, 11, 749.

19 D.C. Appleyard, S.C. Chapin, R.L. Srinivas, P.S. Doyle, Nature Protoc. 2011, 6, 1761.

20 W. Lee, D. Choi, J. Kim, W. Koh, Biomed Microdevices 2008, 10, 813–822.

21 N. Wen, Z. Zhao, B. Fan, D. Chen, D. Men, J. Wang, J. Chen, Molecules 2016, 21, 881.

22 T. Mahmood, P.C. Yang, N. Am. J. Med. Sci. 2012, 4, 429.

23 E. Kinoshita, E. Kinoshita-Kikuta, T. Koike, Nature Protoc. 2009, 4, 1513.

24 M. Butterman, D. Tietz, L. Orbán, A. Chrambach, Electrophoresis 1998, 9, 293–298.

25 J. Tong, J.L. Anderson, Biophys. J. 1996, 70, 1505.

26 J.V. Vlassakis, A.E. Herr, Anal. Chem. 2015, 87, 11030–11038.

27 R.F. Probstein Physicochemical hydrodynamics: an introduction, John Wiley & Sons, 2005.

28 R.J. Kimmerling, G.L. Szeto, J.W. Li, A.S. Genshaft, S.W. Kazer, K.R. Payer, J. de Riba Borrajo, P.C. Blainey, D.J. Irvine, A.K. Shalek, S.R. Manalis Nat. Comm. 2016, 7, 10220.

29 J.P. Beech, S.H. Holm, K. Adolfsson, J.O. Tegenfeldt, Lab Chip 2012, 12, 1048–1051.

30 S.S. Waje, M.W. Meshram, V. Chaudhary, R. Pandey, P.A. Mahanawar, B.N. Braz. J. Chem. Eng. 2005, 22, 209–216.

31 M. Tokita, S. Suzuki, K. Miyamoto, T. Komai, J. Phys. Soc. Jpn. 1999, 68, 330–333.

32 M. Tokita, K. Miyamoto, T. Komai, J. Chem. Phys. 2000, 113, 1647–1650.

33 A.H. Caster, R.A. Kahn, Cell Logistics 2012, 2, 176–188.

34 D. Nguyen, D. Taylor, K. Qian, N. Norouzi, J. Rasmussen, S. Botzet, M. Lehmann, K. Halverson, M. Khine, Lab Chip 2010, 10, 1623–1626.

35 P. Zhu, H.G. Craighead, Annu. Rev. Biophys. 2012, 41, 269–293.

36 J.O. Carvalho, L.P. Silva, R. Sartori, M.A. Dode, PloS one 2013, 8, e59387.

37 H. Sharma, J.B. Wood, S. Lin, R.M. Corn, M. Khine, Langmuir 2014, 30, 10979–10983.

38 K.A. Yamauchi, A.E. Herr, Microsys. Nanoeng. 2017, 3, 16079.

39 B. Gumuscu, J.G. Bomer, A. van den Berg, J.C.T. Eijkel, Biomacromolecules 2015, 16, 3802–3810.

40 D.C. Pregibon, M. Toner, P.S. Doyle, Science 2007, 315, 1393–1396.

